# Plasma-derived extracellular vesicles as potential biomarkers and mediators of functional alterations in MELAS

**DOI:** 10.64898/2026.06.12.731704

**Authors:** Tamiris de Fatima Goebel de Souza, Nicholas Klassen, Patience O. Obi, Berkay Özerkliğ, Tiana Tiede, Abhay Srivastava, Christopher D. Pascoe, Samantha Marin, Sanjiv Dhingra, Cheryl Rockman-Greenberg, Ayesha Saleem

## Abstract

Mitochondrial encephalomyopathy, lactic acidosis and stroke-like episodes (MELAS) syndrome is a genetic disorder characterized by progressive neuromuscular and multisystem symptoms. MELAS typically manifests during childhood, can be difficult to diagnose, and has no cure. Extracellular vesicles (EVs) are lipid-enclosed nanoparticles secreted from cells that contain biological cargo and have demonstrated potential as biomarkers. We investigated the potential of plasma-derived EVs as diagnostic biomarkers of MELAS and examined their functional effects on mitochondrial respiration in treated skeletal muscle myotubes. Plasma-derived EVs were isolated from MELAS patients and age- and sex-matched control individuals, and biophysical characteristics and cargo of EVs analyzed. A Mito Stress Test was performed to assess oxygen consumption rate (OCR) in healthy myotubes treated with Control- or MELAS-EVs to determine the functional effects of circulatory EVs. Nine MELAS patients from two families were studied, and the results were categorized by age, sex and mtDNA heteroplasmy level. EV size and zeta potential remained unchanged. However, total EV concentration was higher in MELAS patients, particularly for small-EVs (<200 nm) and in younger patients (<25 years old). Relative protein yield per EV was lower in the MELAS group, especially among female and younger individuals. EV double-stranded DNA (dsDNA) concentration did not differ between MELAS- and Control-EVs overall, but was higher in male MELAS patients. Protein markers typically enriched in small-EVs showed altered expression in MELAS EVs: TSG101 and CD63 were lower, while flotillin-1 was higher compared to Control-EVs. A decrease in basal OCR was shown in cells treated with MELAS-EVs, with a similar response noted in the group treated with EVs from female MELAS patients. Post-treatment analysis showed no differences in oxidative phosphorylation (OXPHOS) subunit levels between cells treated with MELAS- and Control-EVs. In conclusion, plasma-derived EVs show promise as potential biomarkers for MELAS, and circulating EVs in this patient population may contribute to systemic metabolic dysfunction.

## Introduction

Mitochondrial encephalomyopathy, lactic acidosis, and stroke-like episodes (MELAS) syndrome is a distinctive mitochondrial disorder marked by a wide variety of systemic manifestations (Na and Lee et al., 2024). It is caused by point mutations in mitochondrial DNA (mtDNA), with approximately 80% of cases attributed to the m.3243A>G mutation in the MT-TL1 gene, which encodes for tRNA (Leu-UUR) (Pek et al., 2019). Symptoms typically arise in metabolically active tissues such as skeletal, cardiac muscle, and nervous, which are highly reliant on ATP production, with onset usually occurring before 20 years of age (Thambisetty & Newman, 2014; Tetsuka et al., 2021). However, clinical features can involve nearly all organ symptoms, presenting various combinations, severities, and ages of onset. The heterogeneity in presentation often results in low clinical suspicion, and patients who present with stroke-like episodes, the hallmark symptom of MELAS, are frequently misdiagnosed with acute ischemic stroke (Khasminsky et al., 2023). Current diagnostic methods for MELAS need a combination of multiple approaches, including clinical evaluation, genetic testing, brain imaging, biochemical analysis of blood, cerebrospinal fluid (CSF) and urine, family history, histopathological findings, cardiac and endocrine assessments (Xu et al., 2024). This long and arduous process leads to many patients going years without receiving a diagnosis. This long and complex process makes many patients go undiagnosed for years, emphasizing the importance of biomarkers for early identification to ensure appropriate care.

Extracellular vesicles (EVs) are lipid-bilayer-bound nanoparticles secreted from a variety of cell types and play an important role in cell-to-cell communication. EVs are found in all biological fluids, including blood, saliva, urine, cerebrospinal fluid, amniotic fluid, and more (Welsh et al., 2024). EVs contain biological cargo such as DNA, RNA, lipids, proteins, and metabolites, and have even been shown to contain mitochondrial components and whole mitochondria (Liang et al., 2023). Recent studies indicate that EVs, both small-EVs, (<200 nm) or large-EVs (>200 nm), transport mitochondrial components, including mtDNA, mtRNA, and mitochondrial proteins, that influence cellular and mitochondrial homeostasis (Wang et al., 2025). EVs have been studied as potential biomarkers for many diseases and conditions, such as metabolic syndrome (Martinez & Andriantsitohaina, 2017), autoimmune diseases (Xu et al., 2020), and various types of cancers (An et al., 2015). Notably, damaged-mitochondria enriched EVs are involved in the regulation of several mitochondrial dysfunction-related diseases and may serve as biomarkers for those conditions (Li et al., 2024). One study showed the detection of mtDNA mutations in circulating mitochondrial-derived EVs for pancreatic adenocarcinoma (Vikramdeo et al., 2022). Given that EVs are easily accessible within blood, and the informative profile they provide of the intracellular environment (de Jong et al., 2012), EVs present as putative candidates for biomarkers of diseases with vague clinical presentations such as MELAS syndrome.

To our knowledge, no one has investigated circulating EVs as potential biomarkers of MELAS, nor evaluated the functional effects of these circulatory EVs on perpetuating mitochondrial dysfunction. In this study, we isolated plasma-EVs from MELAS patients along with age- and sex-matched controls, and comprehensively compared EV biophysical characteristics, cargo, and functional effects. We hypothesized that there would be a distinct difference in EV profiles between plasma-EVs from MELAS patients and their respective controls. Additionally, we hypothesized that MELAS-EVs would have a negative functional effect on aerobic metabolism within muscle cells, that may be of clinical relevance for future research into treatments of mitochondrial disorders.

## Methods

### Study participants and ethics approval

The study was designed as a case-control investigation aimed at characterizing plasma-derived EVs from patients with MELAS. The protocol was approved by the ethics committees of the University of Manitoba (under REB# [HS021:341]). Before enrollment, all participants gave written and informed consent. Participant recruitment was coordinated by Dr. Cheryl Rockmann-Greenberg at the Children’s Hospital of Manitoba. Blood samples were collected from 9 participants affected by MELAS and 9 healthy matched controls with no history of mitochondrial disorders or other chronic diseases. Age and sex were used as matching criteria and are shown in Table 1. Patients from two families were included in this study, with the pedigree shown in Fig. 1 and Fig. 2. Heteroplasmy levels in MELAS patients were determined by analysis of peripheral blood samples and reported by the physician. Patient symptoms or signs reported by the physician are detailed in Table 1.

**Figure 1.**
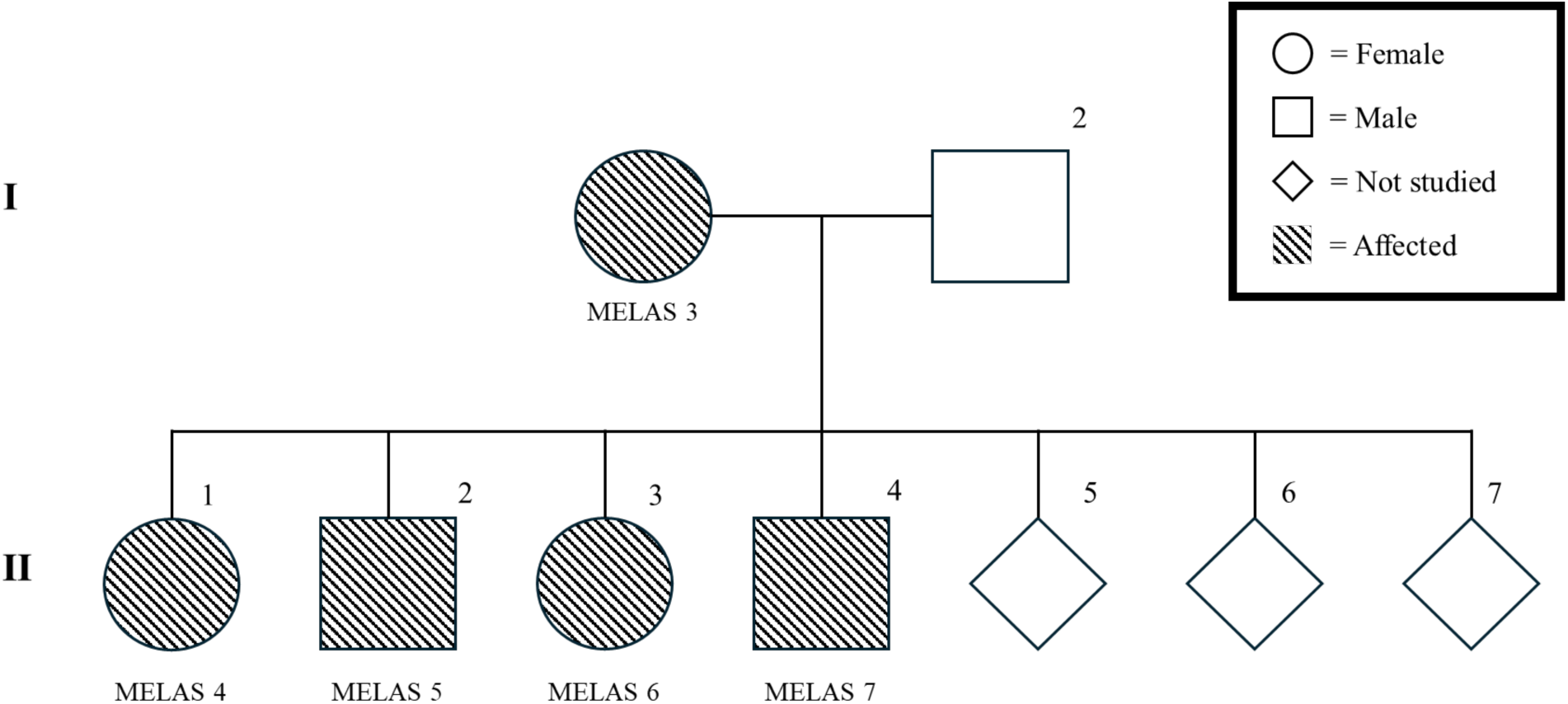
– Family #1 pedigree shows the maternal inheritance of MELAS across two generations. The family tree shows I-1 as a carrier of the m.3234A>G mutation on the mtDNA, along with four descendants (II-1, II-2, II-3 and II-4) with different levels of heteroplasmy. II-1 is the proband of this family.

**Figure 2.**
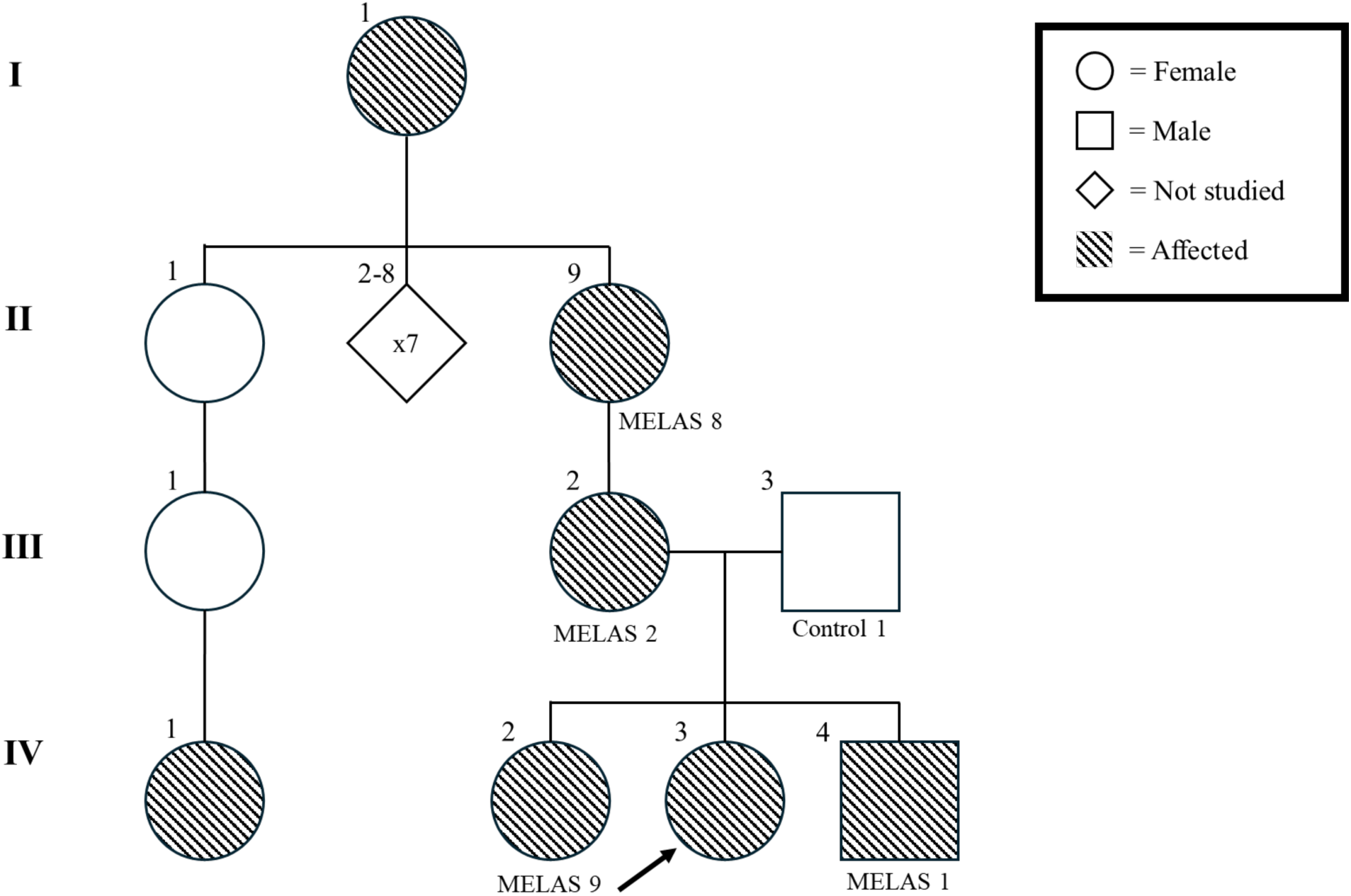
– Family #2 pedigree shows maternal inheritance of MELAS across three generations in a family. The grandmother (II-9) as a carrier of the m.3234A>G mutation on the mtDNA, along with her daughter (III-2) and her three grandchildren (IV-2, IV-3 and IV-4). The father (III-3) was used as a Control for his son (IV-4). IV-4 is the proband of this family.

**Table 1.** Table showing individual characteristics of MELAS subjects from family #1 and family #2. Heteroplasmy was obtained from peripheral blood samples. All controls were asymptomatic.

| **Family #1** |  |  |  |  |  |  |  |  |
| --- | --- | --- | --- | --- | --- | --- | --- | --- |
| Subject | Sex | Birth year | Pedigree # | Subject | Sex | Birth year | Heteroplasmy | Symptoms/signs |
| Control 3 | F | 1991 | I-1 | MELAS 3 | F | 1985 | 0.50% | Asymptomatic (thin with occasional headache) |
| Control 4 | F | 2005 | II-1 | MELAS 4 | F | 2006 | 19.70% | Weight loss, gastroparesis, migraines, epilepsy, psychogenic signs, basal ganglia calcification and stroke |
| Control 5 | M | 2003 | II-2 | MELAS 5 | M | 2007 | 27.30% | Seizures and exercise intolerance |
| Control 6 | F | 2005 | II-3 | MELAS 6 | F | 2010 | 44.40% | Migraines and constipation |
| Control 7 | F | 2016 | II-4 | MELAS 7 | M | 2016 | 61.80% | Asymptomatic |
| **Family #2** |  |  |  |  |  |  |  |  |
| Subject | Sex | Birth year | Pedigree # | Subject | Sex | Birth year | Heteroplasmy | Symptoms/signs |
| Control 1 | M | 1983 | IV-4 | MELAS 1 | M | 2002 | 47.10% | Lactic acidosis, renal failure from interstitial renal disease and febrile seizure at age 2 |
| Control 2 | F | 1982 | III-2 | MELAS 2 | F | 1979 | 18.00% | Short stature, constipation, migraines and psychogenic signs |
| Control 8 | F | 1941 | II-9 | MELAS 8 | F | 1949 | 46.00% | Type 2 diabetes |
| Control 9 | F | 1999 | IV-2 | MELAS 9 | F | 2000 | 16.10% | Migraines, psychogenic signs, constipation, facial tics and abdominal discomfort |

### Plasma collection

Peripheral blood was collected from healthy volunteers (Control, N=9) and affected patients (MELAS, N=9) by venipuncture in the median cubital vein using 21G needles (BD) into 4 mL plasma tubes with dipotassium ethylenediaminetetraacetic acid (K2-EDTA; BD). Tubes were placed on the benchtop for 30 minutes and were centrifuged at 2,000 x g for 10 minutes at room temperature. The upper plasma fraction was separated from blood cells and transferred to 2 mL microcentrifuge tubes in individual aliquots of 110 µL to avoid freeze-thaw cycles. Aliquots were moved to long-term storage at -80 °C.

### Isolation of plasma-derived EVs by size exclusion chromatography

EVs from plasma were isolated using Sepharose-based qEV 35 nm single columns (Izon, New Zealand) according to the manufacturer’s instructions. Plasma aliquot was thawed, and 100 µL of plasma was added to 50 µL of 1X PBS and passed through a 0.2 µm Sartorius Minisart^TM^ RC Syringe Filter (Cat#14555330, Sartorius, Germany). Briefly, the columns were drained and washed with 1X PBS to remove the preservatives. 150 µL of filtered plasma was added to the column and fractions 7-9 (170 µL each) were collected. The eluate was added to Sartorius Vivaspin™ Turbo 4 Centrifugal Concentrator Polyethersulferone, with 10 kDa MWCO (Cat#14-558-708, ThermoFisher, USA) and centrifuged at 7,500 x g for 10 minutes at 4 °C using Sorvall^TM^ RC 6 Plus Centrifuge F13-14 Fixed Angle Rotor (Cat#46922, ThermoFisher, USA). The final volume obtained in the concentrator was collected and transferred to a new 2 mL microcentrifuge tube and used for EV analysis or functional assay.

### Biophysical characterization of EVs by Tunable Resistive Pulse Sensing

Size, concentration, and zeta potential of the plasma EVs were determined by Tunable Resistive Pulse Sensing (TRPS) using the Exoid from Izon (New Zealand). The magnitude of the blockades is directly proportional to the particle diameter, while the blockade rate reflects the particle concentration (Vogel, 2017). Zeta potential is the electric potential difference between the dispersion medium and the layer of fluid attached to the particle, and reflects the stability of a particle in a colloidal suspension. Samples were diluted in a range of 10 to 20 times in a measurement electrolyte (ME) buffer provided by Izon, and were measured using a nanopore 150 (NP150, Izon), which is suitable for analyzing nanoparticles with sizes between 70 and 420 nm. To measure simultaneous size and concentration, each paired sample was analyzed using the same parameters of stretch (46-48 nm), voltage (550-700 mV), and three different pressures (range of 300-1200 Pa) to detect 500 blockades. These settings were adjusted before recording each pair of samples to obtain a current close to 100 nA before beginning the analysis. After sample analysis, carboxylated polystyrene beads (CPC200, Izon) with a known diameter (mean of 200 nm) and stock concentration (8.2E+11 particles/mL) were diluted at 1:750 in ME and used as calibration particles. The calibration particles were analyzed following sample analysis under the same conditions as the sample pairs. For size and zeta analysis, parameters were adjusted to obtain a current close to 130 nA, and 250 blockades were measured at 4 conditions of varying voltages and pressures. Measurements were recorded by the Exoid Control Suite Software (version V1.4.0.88). All the data was analyzed by Exoid Data Suite Software (version V1.0.2.32) to calculate nanoparticle size, concentration, and zeta potential (Lefebvre et al., 2023).

### Protein quantification of plasma EVs

Plasma EVs were lysed using 1:1 lysis buffer with Pierce^TM^ RIPA Solution (Cat#89900, ThermoFisher, USA) with Roche Protease Inhibitor Tablet (Cat#5892791001, Sigma-Aldrich, USA). Samples were vortexed for 5 seconds, followed by two freeze/thaw cycles in liquid nitrogen, then sonicated (30 Hz, 3 x 3 seconds on ice) and centrifuged (15,000 rpm, 15 minutes at 4 °C). The supernatant was collected and used to determine the protein content using the Thermo Scientific TM MicroBCA^TM^ Protein Assay Kit (ThermoFisher, USA) following the manufacturer’s instructions. Briefly, 150 µL working reagent was added to 150 µL of sample or standards (prepared using bovine serum albumin; BSA), in a 96-well plate, incubated for 2 hours at 37°C, and the absorbance was measured at 562 nm using a BioTek Synergy H1 Plate Reader (Agilent, USA). Sample absorbances were compared against the known standard concentrations to determine the protein quantification (µg/mL) of EV lysates (Pierdona et al., 2022).

### EV protein markers quantification by Western blotting

Plasma EV lysates were denatured by the addition of β-mercaptoethanol and 4X Laemmli buffer, followed by incubation at 95 °C for 5 minutes. Equal amounts of protein (2.5 to 5 µg) were loaded onto 12% or 15% SDS-PAGE gels and immersed in 1X running buffer. Gels were run at 90 V for 30 minutes and then 120 V for 1.5 hours, then transferred to a nitrocellulose membrane using Trans-Blot Pierce^TM^ Turbo (Cat#1704150EDU, Bio-Rad Laboratories). The membranes were incubated with Ponceau S Staining Solution (Cat#A40000279, ThermoFisher, USA) for 5 minutes to verify transfer, and the gels were incubated for 15 minutes in Coomassie Blue solution (Cat#1610400, Bio-Rad Laboratories, USA), followed by overnight incubation in destaining solution before imaging. The staining solution was removed by washing for 1 minute in ultrapure water, and membranes were blocked for 5 minutes at room temperature using EveryBlot Blocking Buffer (Cat#12010020, Bio-Rad Laboratories, USA). The membranes were washed 3 x 5 minutes in Tris-buffer saline Tween-20 solution (TBST), then cut based on the size of the target proteins. The cut membranes were incubated overnight at 4 °C with the following primary antibodies diluted in 1% skim milk TBST solution: rabbit polyclonal anti-CD63 (Cat# SAB4301607, Sigma-Aldrich, 1:200), rabbit polyclonal anti-TSG101 (Cat#t5701, Sigma-Aldrich, 1:200), rabbit polyclonal anti-Flotillin-1 (Cat#F1180, Sigma-Aldrich, USA, 1:200), rabbit polyclonal anti-Cytochrome C (Cat#AHP2302, Bio-Rad Laboratories, USA, 1:200), rabbit polyclonal anti-Calnexin (Cat#ab22595, Abcam, USA, 1:1,000), mouse monoclonal anti-heat shock protein (HSP70; Cat#H5147, Sigma-Aldrich, USA, 1:200), mouse monoclonal anti-matrix metalloprotease-2 (MMP-2; Cat#sc-13595, Santa Cruz Biotechnology, USA, 1:200) and mouse monoclonal anti-Apolipoprotein A1 (Apo-A1; Cat#0650-0050, Bio-Rad Laboratories, 1:200). Then, membranes were washed with TBST 3 x 5 minutes and incubated with anti-rabbit IgG secondary antibody (Cat#A16035, ThermoFisher, USA, 1:5,000 or 1:10,000) or anti-mouse IgG secondary antibody (Cat#A16017, ThermoFisher, 1:10,000) diluted in 1% skim milk TBST solution for 1 hour at room temperature. The membranes were washed once more with TBST 3 x 5 minutes before imaging. Membranes were stripped and reprobed by incubating in Thermo Scientific^TM^ Restore^TM^ Western Blot Stripping Buffer (Cat#PI2059, ThermoFisher, USA) for 30 minutes before washing with TBST 3 x 5 minutes and re-blocking the membrane. Clarity Western ECL Substrate (Cat#1705061, Bio-Rad Laboratories, USA) was applied to enhance chemiluminescence and visualize the protein bands. ChemiDoc^TM^ MP Imaging System (Cat#12003154, Bio-Rad Laboratories, USA) was used to acquire images. The images were analyzed using Image Lab software (Bio-Rad Laboratories). Coomassie blue colorimetric images were used as a loading control.

### EV DNA isolation and quantification

EVs were isolated from 100 µL of plasma from each patient and control using qEV columns (as described above). DNA was isolated using the QIAamp DNA Micro Kit (Cat#56304, Qiagen, Germany) using a modified version of the manufacturer’s instructions. The entire concentrated stock of EVs was transferred from the concentrators to a 1.5 mL microcentrifuge tube and the volume was completed to 100 µL with Buffer ATL from the QIAamp DNA Micro Kit. 10 µL of proteinase K, 100 µL of Buffer AL, and 1 µL of RNA carrier (1 µg/µL) was added to the sample and was pulse-vortexed for 15 seconds. The samples were incubated for 10 minutes at 56 °C and sonicated (30 Hz, 3×3 seconds on ice). 50 µL of 100% ethanol anhydrous was added then placed on a rotator mixer for 3 minutes follow by incubation at rest for 3 minutes. The EV lysates were transferred to the QIAamp MiniElute column which was placed inside a 2 mL collection tube and centrifuged at 6,000 x g for 1 minute at room temperature (22 °C). The MiniElute column was then transferred to a clean 2 mL collection tube and 500 µL of Buffer AW1 was added to the column and centrifuged at 6,000 x g for 1 minute at room temperature. This process was repeated once more with 500 µL of Buffer AW2, and centrifuged at 20,000 x g for 3 minutes to dry the membrane. The column was placed in a 1.5 mL collection tube and 20 µL of Buffer AE was added to the centre of the MiniElute column membrane and incubated at room temperature for 5 minutes before being centrifuged at 20,000 x g for 1 minute at room temperature. EV dsDNA was quantified using the Qubit^TM^ dsDNA High Sensitivity (HS) Assay kit (Cat#Q32854, ThermoFisher, USA) and the Qubit^TM^ 4 Fluorometer (Cat#Q33238, ThermoFisher) according to the manufacturer’s instructions. Briefly, 20 µL of isolated DNA from each sample was mixed with 180 µL of Working Solution, vortexed for 3 seconds, and incubated for 4 minutes at room temperature before a reading was taken. Two additional readings were taken at 8 and 12 minutes. A sample of PBS and 1 µL of the RNA carrier was prepared and read in the same manner and subtracted from the final values of sample to control for fluorescence resulting from the RNA carrier.

### C2C12 myotubes differentiation and plasma-EV treatment

Murine C2C12 myoblasts were seeded on a 24-well Seahorse XF Cell Culture Microplate (Cat#100777-004, Agilent, USA) at a density of 3.0E+4 cells/well and were grown in a 37 °C incubator with 5% CO2. Myoblast growth media contained 10% Gibco^TM^ Fetal Bovine Serum (FBS; Cat#A2720801, ThermoFisher, USA) and 1% penicillin/streptomycin (P/S) in Dulbecco’s Modification of Eagle’s Medium (DMEM; Cat# D5796-6X500ML, Sigma-Aldrich, USA). After 24 hours, the media was switched to differentiation media for 5 days to differentiate myoblasts into myotubes. Differentiation media, contained 5% heat-inactivated horse serum (HI-HS; ThermoFisher, USA) and 1% P/S, were refreshed twice during the differentiation process. After C2C12 myotubes completed differentiation, the media were changed to EV-depleted differentiation media (EVd-DM; DMEM supplemented with 5% EV-depleted HI-HS and 1% P/S) for all experimental conditions. The myotubes were treated with fresh plasma EVs isolated from Control or MELAS patients, added in EVd-DM, once a day for two consecutive days. An aliquot of 100 µL of plasma per patient was used to isolate EVs using qEV columns (as described above), and the total yield of EVs was used to treat three wells (three technical replicates per patient). PBS was used as a negative control for EV treatment.

### Mito Stress Test in EV-treated myotubes

After treatment was completed, the media was removed, and cells were washed with XF DMEM assay media, pH 7.4 (Cat#103680-100, Agilent, USA), which was made up of 1% each of: 100 mM pyruvate, 1.0 M glucose, and 200 mM glutamine. Cells were incubated in XF assay media for 60 minutes in a 37 °C, CO2-free incubator. The Seahorse XF Cell Mito Stress Test (Cat#103015-100, Agilent, USA) was performed using the Seahorse XFe24 Analyzer (Agilent, USA) following the manufacturer’s instructions. The key parameters of mitochondrial function are measured through the oxygen consumption rate (OCR) of cells in real time before and after treatment with modulators of key respiratory proteins in the plate well during the assay. Briefly, 1.0 µM oligomycin (complex V inhibitor; Cat#495455, Sigma-Aldrich, USA), 2.0 µM carbonyl cyanide-4-(trifluoromethoxy)-phenylhydrazone (FCCP; uncoupling agent; Cat#C2920, Sigma-Aldrich, USA), 0.5 µM rotenone (complex I inhibitor; Cat#R8875, Sigma-Aldrich, USA), and 0.5 µM antimycin-A (complex III inhibitor; Cat#A8674, Sigma-Aldrich, USA) suspended in XF DMEM assay media were loaded into three different ports of the extracellular flux cartridge. Myotubes were exposed to the modulators in sequential order, and three measurements of OCR taken after each injection, for a total of twelve measurements (three taken before the first injection). Basal and maximal respiration were automatically calculated from these measurements.

### Protein quantification in EV-treated myotubes

Following the Seahorse XF Mito Stress Test, the assay media were disposed of, and the plates were stored upside down in -20 °C overnight. The following day, the plates were thawed, and 50 µL of RIPA lysis buffer was added to each well and incubated for 2 minutes. Each well was then scraped with a 200 µL polystyrene pipette tip and the supernatant was collected and vortexed for 5 seconds followed by two freeze/thaw cycles in liquid nitrogen, then sonicated in a water bath (3 x 2 minutes), and centrifuged (15,000 rpm, 15 minutes at 4 °C). The supernatant was collected and used to determine the protein content using the Thermo Scientific^TM^ Pierce^TM^ BCA Protein Assay Kit (ThermoFisher, USA) according to the manufacturer’s instructions and as described above. Briefly, 200 µL of working reagent was added to 25 µL of sample or BSA standard in a 96-well plate, incubated for 30 minutes at 37 °C, and the absorbance was measured at 562 nm using a microplate reader.

### Electron transport chain (ETC) subunit expression analysis in EV-treated myotube lysates

Myotube lysates were denatured by the addition of β-mercaptoethanol and 4X Laemmli buffer. Incubation at 95 °C was skipped to preserve the detection of Complex IV, as per the manufacturer’s recommendations. Equal amounts of protein (5 µg) were loaded onto 12% SDS-PAGE gels and run as per the protocol described above. An OxPhos cocktail (Cat#45-8099, ThermoFisher, USA, 1:250) containing the antibodies for five respiratory chain proteins corresponding to each complex was used. Membranes were incubated with the primary cocktail antibody for 1 hour at room temperature in a 1% skim milk TBST solution. Afterwards, membranes were washed with TBST 3 x 5 minutes and incubated with anti-mouse IgG secondary antibody (1:10,000) in 1% skim milk TBST solution for 1 hour at room temperature. Clarity Western ECL Substrate solution was applied to enhance chemiluminescence and visualize the protein bands. ChemiDoc^TM^ MP Imaging System was used to acquire images, and the images were analyzed using Image Lab software. Glyceraldehyde-3-phosphate dehydrogenase (GAPDH; Cat#2118S, Cell Signalling, USA; 1:2,000 primary, 1:10,000 secondary) was used as a loading control.

### Statistical analyses

Data were tested for normal distribution using the Shapiro-Wilk test, and analyzed using paired one-tailed t-test for sets that were normally distributed, and a Wilcoxon matched-pairs signed rank test for sets that were not normally distributed. Size-distribution histograms were tested for significance by repeated measures two-way ANOVA. Individual data points are plotted, with ± standard error of mean (SEM) shown as applicable (N=6-9). Significance was set at p ≤0.05, with exact p values shown. GraphPad Prism (Version 10.2.3) software was used for all statistical analyses.

## Results

### Patient characteristics

This study included 9 patients previously diagnosed with MELAS and were matched by age and sex with 9 Control subjects, except for two subjects: MELAS 1 (IV-4), was matched by sex, and MELAS 7 (II-4) matched by age only. A summary of patient and matched control characteristics is presented in **Table 1**. The mean age of the MELAS patients (6 females, 3 males) was 29.1 years (± 20.8, N=9), and the control patients (7 females, 2 males) was 30.1 years (± 22.0, N=9). Peripheral blood heteroplasmy levels show a range of 0.5% to 61.80%, with the highest values found in males (i.e., 61.80.% heteroplasmy) than females (46% heteroplasmy). **Fig. 1** shows Family #1 with five patients, including a mother (I-1) and four descendants (II-1 to II-4), all of whom inherited the mtDNA mutation to varying degrees of heteroplasmy (19.7-61.8%), with an interesting decrease in heteroplasmy levels in the older individuals. **Fig. 2** shows Family #2 with four patients across three generations (II-9, III-2, IV-2 and IV-4), all of whom show maternal inheritance of the same mutation. A case of MELAS was reported in the fourth generation (IV-3) with 56.7% heteroplasmy and clinical manifestations of the disorder, such as headaches, migraines, pseudo seizures, diabetes, and Tourette syndrome, but samples from this subject were not included in this study. Additionally, one other case was reported in the first generation but participant was not included in the study (I-1). MELAS participants and matched controls were grouped by sex [female (N=6) and male (N=3)]; age [<25 years old (N=6) and >25 years old (N=3)]; heteroplasmy level [<20% (N=4) and >20% (N=5)] and family groups [#1 (N=5) and #2 (N=4)] for further analyses.

### Patients with MELAS have higher concentration of circulatory EVs

Plasma-derived EVs from MELAS and Controls were isolated and biophysically characterized by size, zeta potential and concentration using TRPS. Average particle diameter was similar between MELAS-EVs (92.54 nm) and Control-EVs (92.87 nm) (**Fig. 3A**), and zeta potential did not differ between groups (-14.86 mV and -14.70 mV, respectively, **Fig. 3B**). EV distribution reflected an increased plasma EV concentration in MELAS, with MELAS-EVs showing a more dispersed range of small particles (<200 nm) sizes than Control-EVs (**Fig. 3C**). Total plasma EV concentration was 1.96-fold higher (N=9, p=0.027) in MELAS (6.29E+10 particles/mL) than Controls (3.20E+10 particles/mL, **Fig. 3D**). Total EV concentration from plasma-EVs was stratified by sex, age and heteroplasmy levels groups to identify strong patterns. Concentration of MELAS-EVs was trending higher in males (N=3, p=0.0621, **Fig. 3E**), and 1.42-fold higher in young participants <25 years (N=6, p=0.0191, **Fig. 3F**) than Control-EVs. Stratification of EV data by heteroplasmy levels did not reveal any significant patterns in MELAS vs. Controls (**Fig. 3G**).

**Figure 3.**
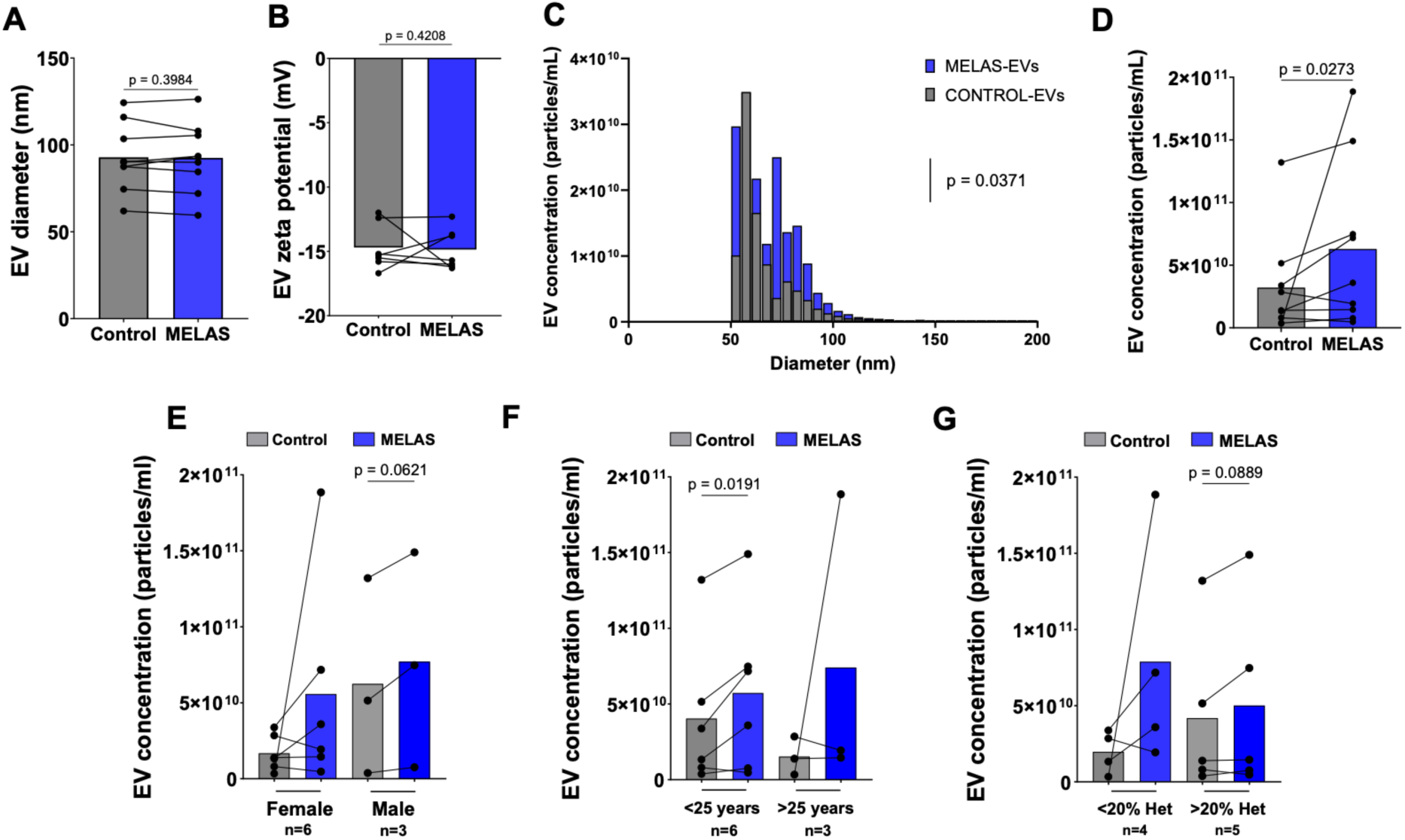
– Concentration of MELAS-EVs is higher in plasma, especially in young individuals. Plasma-derived EVs were isolated via size-exclusion chromatography (SEC) and biophysically characterized by size, zeta potential and concentration using tunable resistive pulse sensing (TRPS). **(A)** The average size (N=9) and **(B)** zeta potential (N=7) remained unchanged between groups. **(C)** The size/concentration distribution histogram representing MELAS and Control small-EVs showed a main effect for MELAS (0-200 nm, N=9, p=0.0371). **(D)** Total EV concentration was 1.97-fold higher in MELAS-EVs (N=9, p=0.0273). **(E)** Stratified data by sex did not show any statistically-significant differences, but trends are noted. **(F)** EV concentration was 1.42-fold higher in <25 years old MELAS individuals (N=6, p=0.0191). **(G)** No differences were observed in EV concentration when stratified by heteroplasmy levels. Each dot on graphs represents one sample, and data are presented as matched control-MELAS pairs. Statistical analyses were performed using a one-tailed paired t-test for data sets that passed the normality test, or Wilcoxon test for those that did not. Normality was assessed using the Shapiro-Wilk test. The histogram was analysed using a two-way ANOVA.

### Sex-specific effects: females with MELAS have lower relative protein content in plasma EVs

Total EV protein yield (µg) was determined by microBCA for all samples (**Fig. 4A**). To estimate the relative EV protein yield (pg/particle), total EV protein yield was normalized by the number of total EV particles in that aliquot (100 µL), as determined by TRPS (**Fig. 4B**). No difference between groups (p=0.4113) was observed for total EV protein yield (**Fig. 4A**). However, there was a 2-fold decrease (N=9, p=0.0020) in protein per particle content in MELAS-EVs (0.012 pg/particle) compared to Control-EVs (0.024 pg/particle) (**Fig. 4B**). The relative protein yield/EV data was stratified by sex, age, heteroplasmy levels and family groups. Protein/EV was 1.95-fold lower in female (N=6, p=0.0368, **Fig. 4C**) and 1.98-fold decreased in <25 years (N=6, p=0.0156, **Fig. 4D**) MELAS subjects. No differences were found when data were stratified by heteroplasmy levels (**Fig. 4E**) or family (**Fig. 4F**) groups.

**Figure 4.**
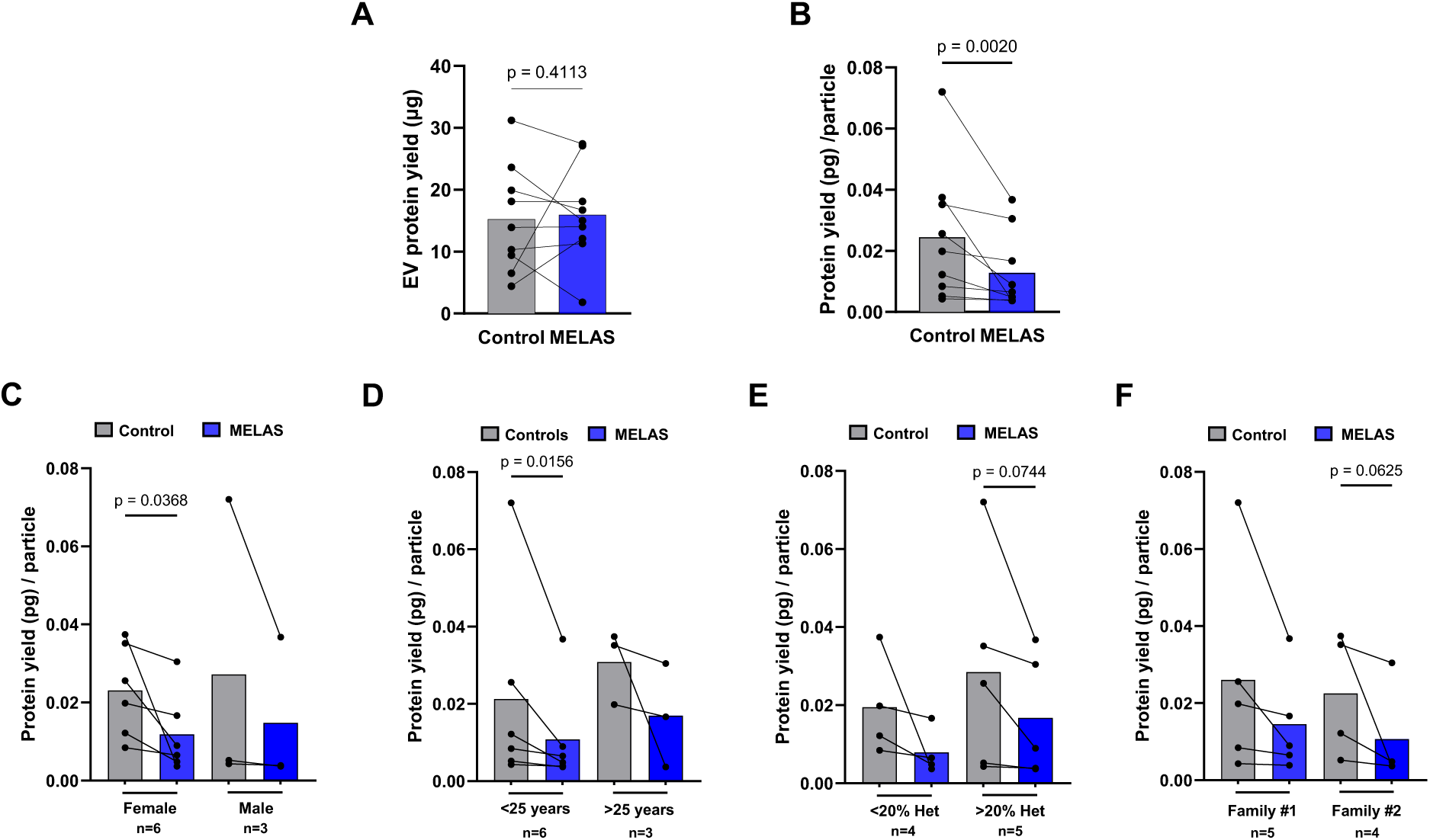
– Relative EV protein yield content is lower in female and young MELAS patients. Plasma-derived EV lysates were used to access EV protein yield using the microBCA assay. Relative protein content was determined by dividing the protein yield (µg) of EV-lysates by the absolute number of particles from same aliquot. **(A)** Total EV protein yield (µg) remained unchanged between groups, but **(B)** relative protein yield (pg/particle) was 2.0-fold lower in MELAS-EVs than Control-EVs (N=9, p=0.0020). **(C)** Relative protein yield/particle was 1.95-fold decreased in females (N=6, p=0.0368), and **(D)** 1.98-fold decreased in <25 years MELAS patients (N=6, p=0.0156). **(E)** Protein yield/particle did not change with heteroplasmy levels, **(F)** nor when stratified by family groups, though notable trends are noted. Each dot on bars represents one sample and data are presented as matched control-MELAS pairs. Statistical analyses were performed using one-tailed paired t-test for data sets that did pass normality test or Wilcoxon test for that did not pass normality according to Shapiro-Wilk test.

### Sex-specific effects: males with MELAS have higher dsDNA content in plasma EVs

Absolute EV dsDNA concentration (ng/mL) was determined (**Fig. 5A**) as described above, and relative dsDNA (pg/particle) determined by normalizing absolute EV dsDNA concentration by EV concentration (**Fig. 5B**). Both absolute and relative dsDNA levels were similar between MELAS and Control-EVs groups. EV ds-DNA concentration data were stratified by sex and heteroplasmy levels groups. Female MELAS-EVs generally had lower absolute dsDNA concentration vs. Controls (N=6, p=0.092), except for one patient (MELAS6, II-3) that was higher than the matched control. Conversely, absolute dsDNA was 2.0-fold higher in male MELAS-EVs than in Control-EVs (N=3, p=0.0027, **Fig. 5C**). Patients with heteroplasmy (<20%) showed generally reduced dsDNA concentration levels (N=4, p=0.0776, **Fig. 5D**), while those with >20% heteroplasmy trended higher absolute dsDNA concentrations (N=5, p=0.3125), except for one patient (MELAS8, II-9), who had lower dsDNA EV-associated than the matched control (**Fig. 5D**).

**Figure 5.**
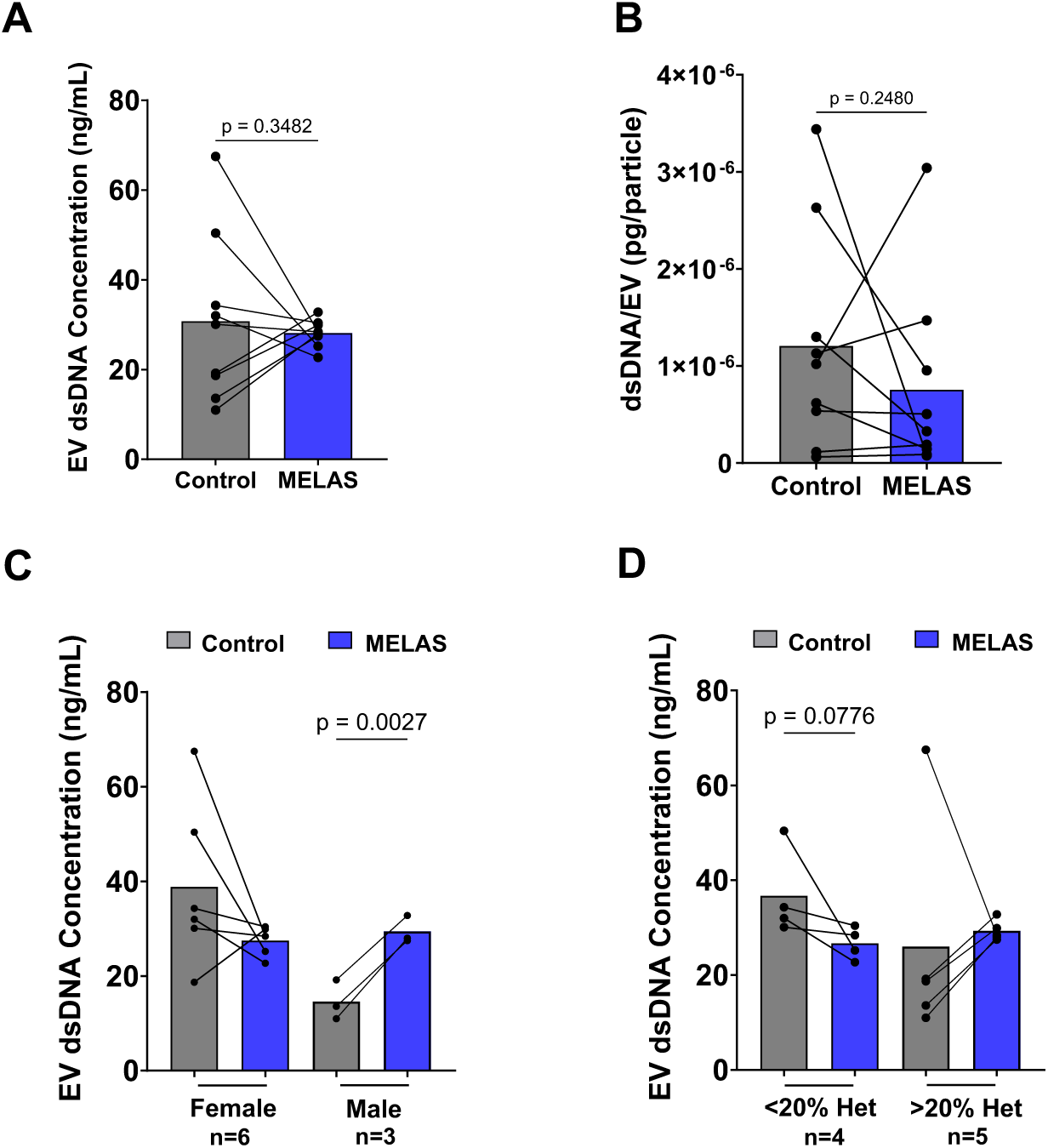
– EV dsDNA content is higher in male MELAS patients. Plasma-derived EV lysates were used to measure the total EV dsDNA concentration (ng/mL). Relative dsDNA (pg/particle) content was determined by dividing the dsDNA yield (µg) of EV-lysates by the absolute number of particles from same aliquot. **(A)** The total EV dsDNA concentration and **(B)** relative dsDNA/EV showed no difference in MELAS-EVs vs. Control-EVs. **(C)** Total EV dsDNA concentration increased 2.0-fold in male MELAS patients (N=3, p=0.0027). **(D)** Stratification of data by heteroplasmy did not show any differences. Each dot on bars represents one sample and data are presented as matched control-MELAS pairs. Statistical analyses were performed using paired t-tests for data sets that passed normality or Wilcoxon tests for data sets that did not pass normality according to Shapiro-Wilk test.

### MELAS-EVs have higher expression of flotillin-1 and lower expression of CD63 and TSG101

We evaluated the purity of our population of EVs by measuring expression of key proteins that are normally enriched in small-EVs (TSG101, flotillin-1 and CD63), along with markers of protein contamination (calnexin). MELAS-EVs showed a 49.8% lower expression of TSG101 (N=8, p=0.0197, **Fig. 6A**) and a 64.5% decreased expression of CD63 (N=9, p<0.0001, **Fig. 6C**). Flotillin-1 expression was 1.90-fold higher in MELAS-EVs compared to Control-EVs (N=9, p=0.0303, **Fig. 6B**). Additionally, all EV-lysates were negative for calnexin, an endoplasmic reticulum marker used to assess cellular contamination (**Fig. 6D**).

**Figure 6.**
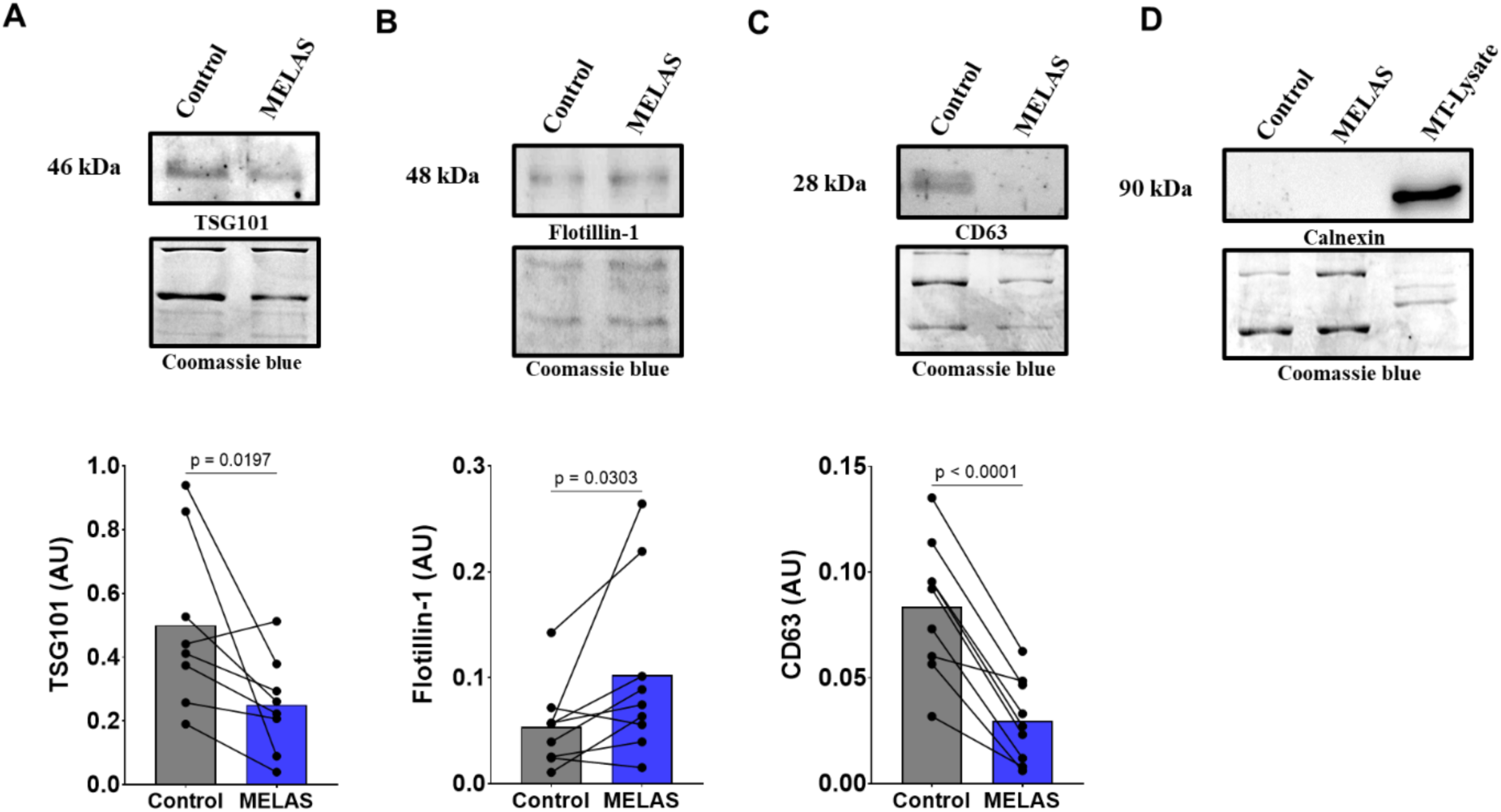
– MELAS-EVs show different expression of EV-proteins markers. Plasma-EV lysates were denatured and equal amount of protein (2.5 to 5 µg) were loaded onto 12% or 15% SDS-PAGE gels for protein separation and western blotting analysis. The images show representative bands for Control and MELAS-EVs illustrated for each protein above quantified results for all samples. (**A**) TSG101 expression was decreased by 49.8% (N=8, p=0.0197), (**B**) flotillin-1 expression was increased by 52.5% (N=9,p=0.0303), and (**C**) CD63 expression was decreased by 64.5% (N=9, p<0.001) in MELAS-EVs vs. controls. All EV lysates were negative for endoplasmic reticulum marker calnexin (**D**). Bands were corrected for loading by Coomassie blue, with multiple proteins run on one gel. Each dot on bars represents one sample and data are presented as matched control-MELAS pairs. Statistical analyses were performed using paired t-test.

### MELAS EV-treated myotubes have decreased basal oxygen consumption rates (OCR)

C2C12 myotubes were cultivated and differentiated in an XFe24-well plate, and treated with plasma-EVs for 2 days, once a day as detailed above. Mito stress assay was performed after 24 hours from the last treatment to access functional effects of plasma-derived MELAS-EVs on healthy cells. The OCR (pmol/min/µg) curve shows the cells response to the exposure of inhibitors after MELAS- and Control-EVs treatment (**Fig. 7A**). Cells treated with MELAS-EVs had a 12.8% decrease in basal OCR (N=9, p=0.0292, **Fig. 7B**), with no significant effect on maximal OCR (N=9, p=0.2195, **Fig. 7C**). Sex-stratified data illustrated lower basal OCR in cells treated with female MELAS EVs vs. Controls (N=6, p=0.0292, **Fig. 7D**). However, no differences were observed in data stratified by age- (**Fig. 7E**) and heteroplasmy levels (**Fig. 7F**).

**Figure 7.**
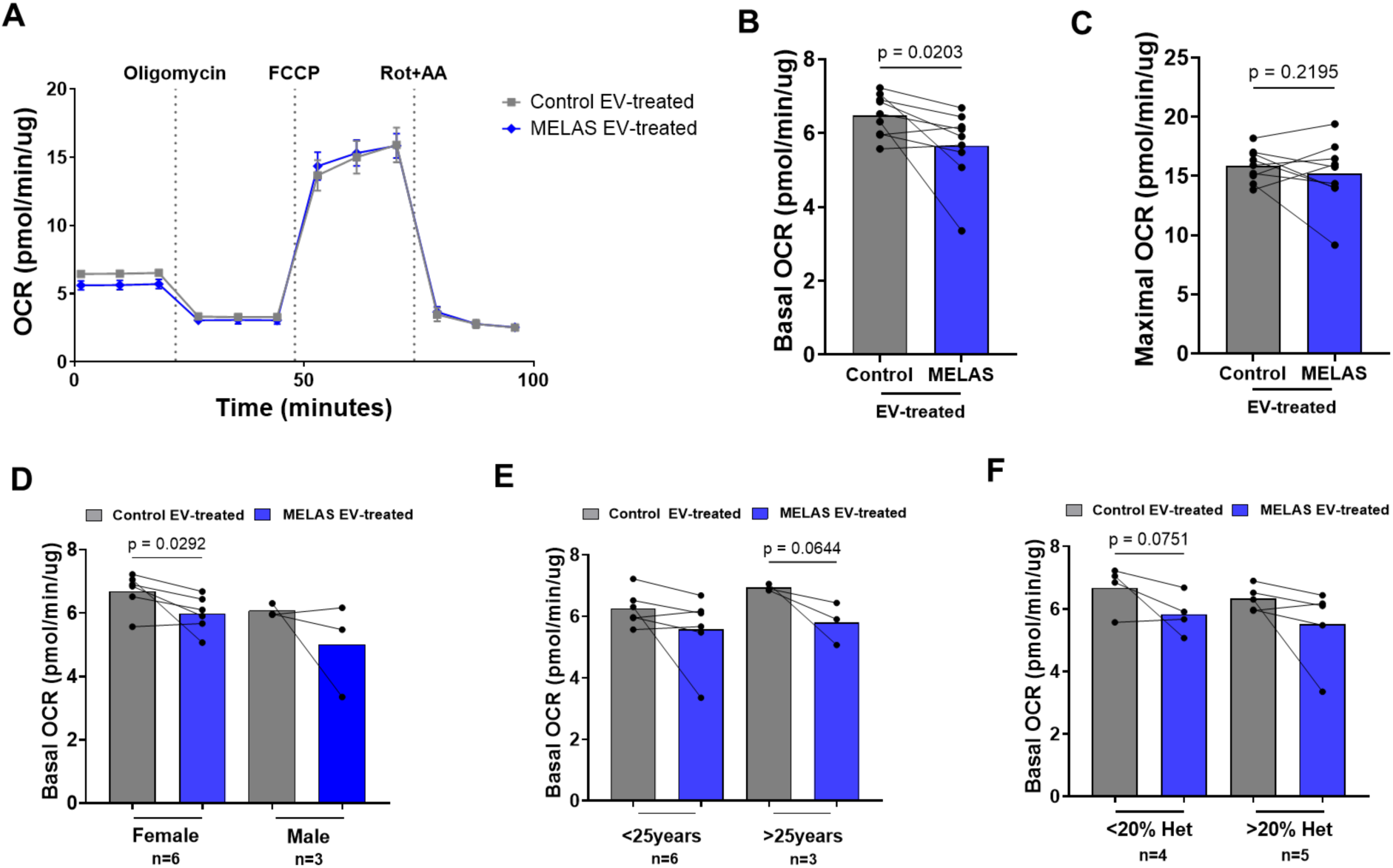
– MELAS-EVs treatment reduced oxygen consumption rates (OCR) in healthy C2C12 myotubes. Mito stress assay was performed 24hours after cells were treated with plasma-EVs from either group (once/day, 2 days). The values were normalized to the total cellular protein per well (µg). (**A**) The OCR (pmol/min/µg) curve shows the cells response to the exposure of oligomycin (1µM), FCCP (2µM), rotenone (0.5 µM) and antimicyn A (0.5 µM) after treatment with plasma-EVs. (**B**) Basal OCR in MELAS EV-treated myotubes were reduced by 12.8% % (N=9, p=0.0203). (**C**) Maximal OCR did not show changes with EV treatment. **(D)** Female MELAS EVs reduced basal OCR (N=6, p=0.0292) vs. Control EV-treated cells, and no significant effects were observed in male MELAS-EV group. **(E)** Age-stratified and **(F)** Heteroplasmy-stratified basal OCR did not show any statistical differences between groups, but notable trends are recorded. Each dot on bars represents one sample and data are presented as matched control-MELAS pairs. Statistical analyses were performed using one-tailed paired t-test.

### Respiratory chain protein expression from myotubes treated with MELAS-EVs does not differ

We evaluated the expression of subunits of protein complexes of the respiratory chain complexes in EV-treated myotubes to determine the effect of MELAS EVs on expression of proteins related to mitochondrial function. While majority of data was not statistically significant (**Fig. 8A-C, Fig. 8E, Fig. 8F**), Complex IV (MTCO1) expression was reduced by 25.7% (N=6, p=0.0817, **Fig. 8D**), in MELAS-treated myotubes compared with Controls.

**Figure 8.**
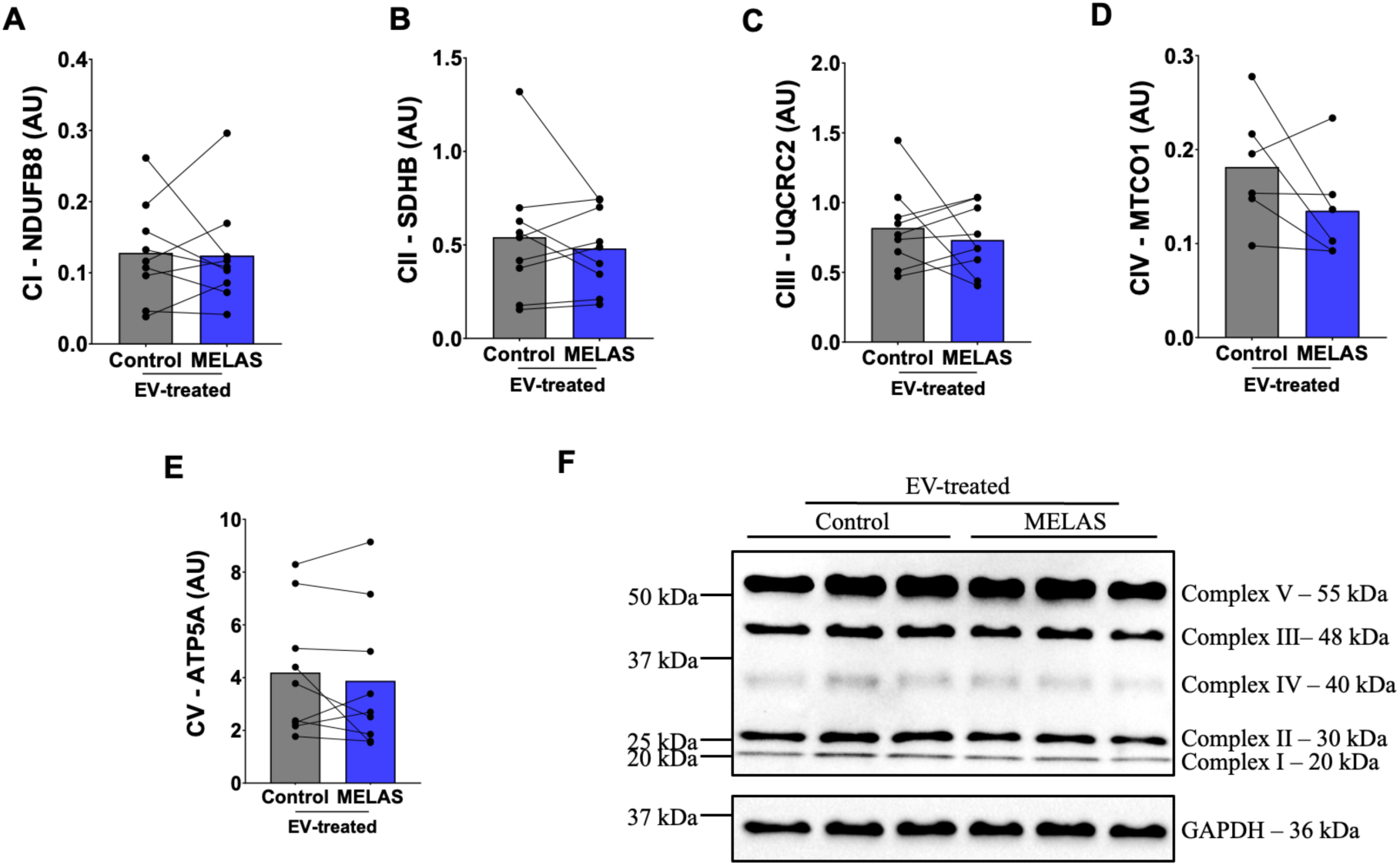
– Expression of proteins subunits from respiratory chain complexes in myotubes treated with plasma-EVs derived from MELAS or Control subjects. (**A**) Complex I: NADH dehydrogenase [ubiquinone] 1 beta subunit subcomplex 8 (NDUFB8) (N=9), (**B**) Complex II: succinate dehydrogenase [ubiquinone] iron-sulfur subunit (SDHB) (N=9), and (**C**) Complex III: cytochrome b-c1 complex subunit 2 (UQCRC2) (N=9) did not show any changes with MELAS-EV treatment. (**D**) Complex IV: mitochondrially encoded cytochrome c oxidase I (MTCO1) (N=6, p=0.0817) showed a trend towards decrease in expression with MELAS-EVs. (**E**) Lastly, no effect of treatment was seen in complex V: ATP synthase, H+ transporting, F1 complex, alpha a (ATP5A) (N=9) expression. (**F**) Representative western blot for OxPhos protein expression is shown. Bands were corrected for loading by glyceraldehyde-3-phosphate dehydrogenase (GAPDH). Each dot on bars represents one sample and data are presented as matched control-MELAS pairs. Statistical analyses were performed using paired t-test for data sets that passed normality or Wilcoxon t-tests for data sets that did not pass normality according to Shapiro-Wilk test.

## Discussion

In this study, we investigated the difference in biophysical characteristics, cargo, and functional effects of plasma-EVs from patients with MELAS. To our knowledge, this is one of the first studies analyzing EVs derived from patients with MELAS. The results of this study support our hypothesis that there is a distinct and salient difference in biophysical properties and biological cargo of plasma-EVs from patients with MELAS, which may be clinically relevant if EVs can serve as putative biomarkers of this hard to diagnose disorder. In particular we found that MELAS patients have an increased concentration of EVs, lower expression of EV proteins TSG101 and CD63, and lower dsDNA EV cargo specifically in male patients. We also investigated the functional effects of plasma-EVs from MELAS patients *in vitro*. Treated myotubes exhibited decreased basal OCR after two days of treatment with MELAS-EVs, suggesting that circulating EVs in these patients may be perpetuating dysfunction in aerobic metabolism. The mechanisms behind the changes in EV characteristics and functional effects in MELAS have yet to be determined.

The increase in plasma-EV concentration in MELAS patients is consistent with previous research in other acute and pathological stressors, such as exercise (Frühbeis et al., 2015), viral and bacterial infections (Chen et al., 2020; Barberis et al., 2021), trauma (Curry et al., 2014), autoimmune diseases (Bartoloni et al., 2015; Burbano et al., 2018), neurodegenerative diseases (Picca et al., 2020a), and various cancers (Baran et al., 2010). There were extreme variations in plasma-EV concentration between patient and control samples, ranging from no difference to a 56-times higher concentration. This had no apparent correlation with the heteroplasmy levels of the patients. It should be noted that the EVs isolated came from the bloodstream, which contains EVs originating from a variety of cell types and due to different biogenesis pathways. Further investigation into specific populations of EVs, such as from skeletal muscle or the central nervous system (Shi et al., 2014), may provide tissues-specific differences in the characteristics of EVs from MELAS patients vs. healthy controls.

Despite a higher concentration of plasma-EVs in MELAS patients, the relative EV protein load was lower in MELAS compared to matched controls. This may indicate a distinct subpopulation of EVs released with protein cargo specificity. Indeed, we measured salient differences in expression of small-EV protein markers (e.g., TSG101, flotillin-1, and CD63) between MELAS and Control EVs. Flotilin-1 is a lipid raft-associated protein that functions as a scaffold for membrane remodeling and plays important roles in cell adhesion, signal transduction, lipid homeostasis and membrane trafficking. Increased flotillin-1 expression has been reported in several pathological conditions, including cancer, neurodegenerative diseases, and metabolic diseases (Bodin et al., 2025). In the present study, flotillin-1 expression was elevated in small-EVs isolated from MELAS patients, suggesting a potential role in the widespread cellular dysfunction characteristic of this mitochondrial disorder.

In contrast, the expression of TSG101 and CD63 was reduced in MELAS-EVs. TSG101 is an abundant cytosolic protein involved in intraluminal vesicle formation and endosomal trafficking and is widely recognized as a canonical small-EV marker (Koritzinsky et al., 2019). Notably, cardiac overexpression of TSG101 has been shown to restore impaired mitophagy in lipopolysaccharide-challenged mouse hearts, resulting in improved cardiac function and survival (Essandoh et al., 2019). Given that defective mitophagy is a hallmark of all mitochondrial disorders caused by pathogenic mtDNA mutations, reduced TSG101 levels in MELAS-EVs may reflect alterations in mitophagy-related processes and endosomal trafficking pathways, potentially contributing to the accumulation of dysfunctional mitochondria and disease pathogenesis. The tetraspanin protein CD63 had lower expression across all samples, implicating its use as a potential biomarker and its likely role in the pathological processes in MELAS syndrome. Previously increased small-EV secretion with lower CD63 expression was reported in the serum of older individuals with physical frailty and sarcopenia, putatively attributed to disarrangements in late endocytic trafficking (Picca et al., 2020b). However, in our study, some patients are young and presented the same pattern as aged individuals, suggesting that the specificity of CD63 expression is likely not due to age, but rather to mitochondrial dysfunction. In addition, previous research has shown varied CD63 expression patterns in EVs, which were increased in some cancers (Odaka et al., 2022; Zhou et al., 2021), while decreased expression was inversely correlated with metastatic potential (Chen et al., 2011; Liu et al., 2018). The role of CD63 in cancer diagnostics and prognostics seems to depend on a variety of factors, exhibiting both anti- and pro-tumorigenic effects depending on the cancer type (Dey et al., 2023). Elucidating the mechanisms underlying this robust decrease in CD63 in MELAS patients irrespective of sex, age, and heteroplasmy levels, is required before it can be used as a putative biomarker. Interestingly, we found that circulatory MELAS-EVs exert a detrimental effect on metabolic capacity in muscle cells, which is a clinically relevant finding for patients. Myotubes treated with plasma-EVs derived from MELAS patients exhibited decreased basal OCR compared to their respective controls. Western blot analysis of respiratory chain proteins revealed no significant difference in expression. However, it is worth noting that the decrease in expression of complex IV subunit with MELAS-EV treatment neared statistical significance, as this protein is encoded by mtDNA. The functional effect on OCR is striking given the short treatment time; a concomitant decrease in the expression of proteins involved in mitochondrial biogenesis pathways may take longer treatment time. Stratification by heteroplasmy into low (under 20%) and high (over 20%) revealed a slight trend towards decreased expression of respiratory chain proteins in the high heteroplasmy group, however, the difference was non-significant. We do not know if the decrease in OCR is due to suppression of canonical pathways involved in mitochondrial biogenesis or through indirect mechanisms such as delivery of iron by EVs as illustrated by Yanatori et al. (2021). Further research should investigate the expression of key iron-regulating proteins, such as ferritin, and iron content in MELAS-EVs.

Our study has a few key limitations. First, our small sample size is derived from only two families, suggesting the possibility of an unrelated genetic explanation for the differences we observed. Second, the heteroplasmy reported was derived from peripheral blood mononuclear cells and is likely different from the heteroplasmy in more affected tissues such as the nervous system and skeletal muscle. Previous research has shown that skeletal muscle heteroplasmy is consistently higher and remains constant over time, while blood heteroplasmy reduces with age (Rajasimha et al., 2008). Further investigation into the correlation between heteroplasmy and symptom severity should consider heteroplasmy in hair follicles and muscle biopsies which may be more sensitive and reliable (Sue et al., 1998). Lastly, our investigation of the functional effects of MELAS-derived EVs in myotubes took place over three days, with two total treatments and used murine cells. A more accurate simulation of the effects of circulating EVs in MELAS patients would involve chronic exposure over a longer period, and using primary human cells.

In summary, we investigated the biophysical characteristics and cargo of EVs derived from patients with MELAS to evaluate their feasibility as a putative blood-based biomarker. We also investigated the functional effects of MELAS-derived EVs to elucidate their role in the pathophysiology of the disorder and highlight any potential avenue for therapeutic interventions. Our findings demonstrated a clear difference in the concentration of EVs in the blood of MELAS patients, along with differential protein expression. We also demonstrated that MELAS-derived EVs decreased metabolic capacity in myotubes, implicating their role in perpetuating metabolic dysfunctions systemically. The underlying mechanisms, and whether these observations are unique to MELAS or applicable to other mitochondrial disorders are not yet known.

## References

An, T., Sihua, Q., Xu, Y., Tang, Y., Huang, Y., Situ, B., Inal, J.M., & Zheng, L. (2015). Exosomes serve as tumour markers for personalized diagnostics owing to their important role in cancer metastasis. Journal of Extracellular Vesicles, 4(1). 10.3402/jev.v4.27522

Baran, J., Baj-Krzyworzeka, M., Weglarczyk, K., Szatanek, R., Zembala, M., Barbasz, J., Szczepanik, A., & Zembala, M. (2010). Circulating tumour-derived microvesicles in plasma of gastric cancer patients. Cancer Immunology, Immunotherapy, 59, 841–850. 10.1007/s00262-009-0808-2

Barberis, E., Vanella, V.V., Falasca, M., Caneapero, V., Cappellano, G., Raineri, D., Ghirimoldi, M., De Giorgis, V., Puricelli, C., Vaschetto, R., Sainaghi, P.P., Bruno, S., Sica, A., Dianzani, U., Rolla, R., Chiocchetti, A., Cantaluppi, V., Baldanzi, G., Marengo, E., & Manfredi, M. (2021). Circulating exosomes are strong involves in SARS-CoV-2 infection. Frontiers in Molecular Bioscience, 8. 10.3389/fmolb.2021.632290

Bartoloni, E., Alunno, A., Bistoni, O., Caterbi, S., Luccioli, F., Santoboni, G., Mirabelli, G., Cannarile, F., & Gerli, R. (2015). Characterization of circulating endothelial microparticles and endothelial progenitor cells in primary Sjögren’s syndrome: New markers of chronic endothelial damage? Rheumatology, 54(3), 536–544. 10.1093/rheumatology/keu320

Bodin, S., Elhabashy, H., Macdonald, E., Winter, D., & Gauthier-Rouvière, C. (2025). Flotillins in membrane trafficking and physiopathology. Biology of the Cell, 117, e2400134. 10.1111/boc.202400134

Burbano, C., Roja, M., Muñoz-Vahos, C., Vanegas-García, A., Correa, L.A., Vásquez, G., & Castaño, D. (2018). Extracellular vesicles are associated with the systemic inflammation of patients with seropositive rheumatoid arthritis. Scientific Reports, 8(1), 17917. 10.1038/s41598-018-36335-x

Chen, Z., Gu, S., Trojanowicz, B., Liu, N., Zhu, G., Dralle, H., & Hoang-Vu, C. (2011). Down-regulation of TM4SF is associated with the metastatic potential of gastric carcinoma TM4SF members in gastric carcinoma. World Journal of Surgical Oncology, 9, 43. 10.1186/1477-7819-9-43

Chen, H.P., Wang, X.Y., Pan, X.Y., Hu, W.W., Cai, S.T., Joshi, K., Deng, L.H., & Ma, D. (2020). Circulating neutrophil-derived microparticles associated with prognosis of patients with sepsis. Journal of Inflammation Research, 13, 1113–1124. 10.2147/JIR.S287256

Cotticelli, M.G., Crabbe, A.M., Wilson, R.B., & Shchepinov, M.S. (2013). Insight into the role of oxidative stress in the pathology of Friedreich ataxia using peroxidation resistant polyunsaturated fatty acids. Redox Biology, 1(1), 398–404. 10.1016/j.redox.2013.06.004

Curry, N., Raja, A., Beavis, J., Stanworth, S., & Harrison, P. (2014). Levels of procoagulant microvesicles are elevated after traumatic injury and platelet microvesicles are negatively correlated with mortality. Journal of Extracellular Vesicles, 3(1), 25625. 10.3402/jev.v3.25625

de Jong, O.G., Verhaar, M.C., Chen, Y., Vader, P., Gremmels, H., Posthuma, G., Schiffelers, R.M., Gucek, M., & van Balkom, B.W.M. (2012). Cellular stress conditions are reflected in the protein and RNA content of endothelial cell-derived exosomes. Journal of Extracellular Vesicles, 1(1), 18396. 10.3402/jev.v1i0.18396

Delatycki, M.B., Camarakis, J., Brooks, H., Evans-Whipp, T., Thorburn, D.R., Williamson, R., & Forrest, S.M. (1999). Direct evidence that mitochondrial iron accumulation occurs in Friedreich ataxia. Annals of Neurology, 45, 673–675.

Delatycki, M.B., & Corben, L.A. (2012). Clinical features of Friedreich Ataxia. Journal of Child Neurology, 29(9), 1133–1137. 10.1177/0883073812448230

Dey, S., Basu, S., & Ranjan, A. (2023). Revisiting the role of CD63 as pro-tumorigenic or anti-tumorigenic tetraspanin in cancers and its theragnostic implications. Advanced Biology, 7(7), 2300078. https://doi-org.uml.idm.oclc.org/10.1002/adbi.202300078

Essandoh K, Wang X, Huang W, Deng S, Gardner G, Mu X, Li Y, Kranias EG, Wang Y, Fan GC. Tumor susceptibility gene 101 ameliorates endotoxin-induced cardiac dysfunction by enhancing Parkin-mediated mitophagy. (2019) J Biol Chem. 294(48):18057–18068. doi: 10.1074/jbc.RA119.008925

Frühbeis, C., Helmig, S., Tug, S., Simon, P., & Krämer-Albers, E. (2015). Physical exercise induces rapid release of small extracellular vesicles into the circulation. Journal of Extracellular Vesicles, 4(1), 28239. 10.3402/jev.v4.28239

Khasminsky, V., Auriel, E., Luckman, J., Eliahou, R., Inbar, E., Pardo, K., Landau, Y, Barnea, R., Mermelstein, M., Shelly, S., Naftali, J., & Peretz, S. (2023). Clinicoradiologic criteria for the diagnosis of stroke-like episodes in MELAS. Neurology Genetics, 9(4). 10.1212/NXG.0000000000200082

Kispal, G., Csere, P., Prohl, C., & Lill, R. (1999). The mitochondrial proteins Atmp1p and Nfs1p are essential for biogenesis of cytosolic Fe/S proteins. The EMBO Journal, 18(14). 3981–3989. 10.1093/emboj/18.14.3981

Koritzinsky EH, Street JM, Chari RR, Glispie DM, Bellomo TR, Aponte AM, Star RA, Yuen PST. Circadian variation in the release of small extracellular vesicles can be normalized by vesicle number or TSG101. (2019). Am J Physiol Renal Physiol., 1(317):F1098–F1110. doi: 10.1152/ajprenal.00568.2017

Lefebvre, A., Trioën, C., Renaud, S., Laine, W., Hennart, B., Bouchez, C., Leroux, B., Allorge, D., Kluza, J., Werkmeister, E., Grolez, G.P., Delhem, N., & Moralès, O. (2023). Extracellular vesicles derived from nasopharyngeal carcinoma induce the emergence of mature regulatory dendritic cells using a galectin-9 dependent mechanism. Journal of Extracellular Vesicles, 12(12), 12390. 10.1002/jev2.12390

Li, J., Wang, T., Hou, X. et al. (2024). Extracellular vesicles: opening up a new perspective for the diagnosis and treatment of mitochondrial dysfunction. J Nanobiotechnol 22, 487. 10.1186/s12951-024-02750-8

Liang, W., Sagar, S., Ravindran, R., Najor, R.H., Quiles, J.M., Chi, L., Diao, R.Y., Woodall, B.P., Leon, L.J., Zumaya, E., Duran, J., Cauvi, D.M., De Maio, A., Adler, E.D., & Gustafsson, Å. B. (2023). Mitochondria are secreted in extracellular vesicles when lysosomal function is impaired. Nature Communications, 14, 5031. 10.1038/s41467-023-40680-5

Lill, R. (2009). Function and biogenesis of iron-sulphur proteins. Nature, 460, 831–838. 10.1038/nature08301

Lin, C.M., & Thajeb, P. (2007). Valproic acid aggravates epilepsy due to MELAS in a patient with an A3243G mutation of mitochondrial DNA. Metabolic Brain Disease, 22, 105–109. 10.1007/s11011-006-9039-9

Liu, W.H., Li, X., Zhu, X.L., Hou, M.L., & Zhao, W. (2018). CD63 inhibits the cell migration and invasion ability of tongue squamous cell carcinoma. Oncology Letters, 15(6), 9033–9042. https://doi-org.uml.idm.oclc.org/10.3892/ol.2018.8499

Martinez, M.C., & Andriantsitohaina, R. (2017). Extracellular vesicles in metabolic syndrome. Circulation Research, 120(10), 1674–1686. 10.1161/CIRCRESAHA.117.309419

Mihaylova, M.M., & Shaw, R.J. (2011). The AMP-activated protein kinase (AMPK) signalling pathway coordinates cell growth, autophagy, & metabolism. Nature Cell Biology, 13(9), 1016–1023. https://doi-org.uml.idm.oclc.org/10.1038/ncb2329

Na JH, Lee YM. (2024). Diagnosis and Management of Mitochondrial Encephalopathy, Lactic Acidosis, and Stroke-like Episodes Syndrome. Biomolecules. 2024 Nov 28;14(12):1524. doi: 10.3390/biom14121524

Odaka, H., Hiemori, K., Shimoda, A., Akiyoshi, K., & Tateno, H. (2022). CD63-positive extracellular vesicles are potential diagnostic biomarkers of pancreatic ductal adenocarcinoma. BMC Gastroenterology, 22, 153. 10.1186/s12876-022-02228-7

Palikaras, K., Lionaki, E., & Tavernarakis, N. (2018). Mechanisms of mitophagy in cellular homeostasis, physiology and pathology. Nature Cell Biology, 20, 1013–1022. 10.1038/s41556-018-0176-2

Pek NMQ, Phua QH, Ho BX, Pang JKS, Hor JH, An O, Yang HH, Yu Y, Fan Y, Ng SY, Soh BS. (2019). Mitochondrial 3243A > G mutation confers pro-atherogenic and pro-inflammatory properties in MELAS iPS derived endothelial cells. Cell Death Dis. 10(11):802. doi: 10.1038/s41419-019-2036-9. PMID: 31641105; PMCID: PMC6805858.

Picca, A., Guerra, F., Calvani, R., Marini, F., Biancolillo, A., Landi, G., Raffaella, B., Landi, F., Bernabei, R., Bentivoglio, A.R., Lo Monaco, M.R., Bucci, C., & Marzetti, E. (2020). Mitochondrial signatures in circulating extracellular vesicles of older adults with Parkinson’s disease: Results from the Exosomes in Parkinson’s Disease (EXPAND) study. Journal of Clinical Medicine, 9(2), 504. 10.3390/jcm9020504

Pierdoná TM, Martin A, Obi PO, Seif S, Bydak B, Labouta HI, Eadie AL, Brunt KR, McGavock JM, Sénéchal M, Saleem A. (2022). Extracellular Vesicles as Predictors of Individual Response to Exercise Training in Youth Living with Obesity. Front Biosci (Landmark Ed). 27(5):143. doi: 10.31083/j.fbl2705143.

Puccio, H., Simon, D., Cossée, M., Criqui-Filipe, P., Tiziano, F., Melki, J., Hindelang, C., Matyas, R., Rustin, P., & Koenig, M. (2001). Mouse models for Friedreich ataxia exhibit cardiomyopathy, sensory nerve defect, and Fe-S enzyme deficiency followed by intramitochondrial iron deposits. Nature Genetics, 27, 181–186. https://doi-org.uml.idm.oclc.org/10.1038/84818

Rajasimha, H.K., Chinnery, P.F., & Samuels, D.C. (2008). Selection against pathogenic mtDNA mutations in a stem cell population leads to the loss of the 3243AwG mutation in blood. American Journal of Human Genetics, 82(2), 333–343. 10.1016/j.ajhg.2007.10.007

Shi, M., Liu, C., Cook, T.J., Bullock, K.M., Zhao, Y., Ginghina, C., Li, Y., Aro, P., Dator, R., He, C., Hipp, M.J., Zabetian, C.P., Peskin, E.R., Hu, S., Quinn, J.F., Galasko, D.R., Banks, W.A., & Zhang, J. (2014). Plasma exosomal α-synuclein is likely CNS-derived and increased in Parkinson’s disease. Acta Neuropathologica, 128, 639–650. https://doi-org.uml.idm.oclc.org/10.1007/s00401-014-1314-y

Shuler, K.T., Wilson, B.E., Muñoz, E.R., Mitchell, A.D., Selsby, J.T., & Hudson, M.B. (2020). Muscle stem cell-derived extracellular vesicles reverse hydrogen peroxide-induced mitochondrial dysfunction in mouse myotubes. Cells, 9(12), 2544. 10.3390/cells9122544

Sproule, D.M., & Kaufmann, P. (2008). Mitochondrial encephalopathy, lactic acidosis, and strokelike episodes. Annals of the New York Academy of Sciences, 1142(1), 133–158. https://doi-org.uml.idm.oclc.org/10.1196/annals.1444.011

Sue, C.M., Quigley, A., Katsabanis, S., Kapsa, R., Crimmins, D.S., Byrne, E., & Morris, J.G.L. (1998). Detection of MELAS A3243G point mutation in muscle, blood and hair follicles. Journal of the Neurological Sciences, 161(1), 36–39. 10.1016/S0022-510X(98)00179-8

Tetsuka S, Ogawa T, Hashimoto R, Kato H. (2021). Clinical features, pathogenesis, and management of stroke-like episodes due to MELAS. Metab Brain Dis. 36(8):2181–2193. doi: 10.1007/s11011-021-00772-x

Thambisetty, M., & Newman, N.J. (2004). Diagnosis and management of MELAS. Expert Review of Molecular Diagnostics, 4(5), 631–644. 10.1586/14737159.4.5.631

Vanderboom, P.M., Dasari, S., Ruegsegger, G.N., Pataky, M.W., Lucien, F., Heppelmann, C.J., Lanza, I.R., & Nair, K.S. (2021) A size-exclusion-based approach for purifying extracellular vesicles from human plasma. Cell Reports Methods, 1(3), 100055. 10.1016/j.crmeth.2021.100055

Vashisht, A.A., Zumbrennen, K.B., Huang, X., Powers, D.N., Durazo, A., Sun, D., Bhaskaran, N., Persson, A., Uhlen, M., Sangfelt, O., Spruck, C., Leibold, E.A., & Wohlschlegel, J.A. (2009). Control of iron homeostasis by an iron-regulated ubiquitin ligase. Science, 326(5953), 718–721. https://doi-org.uml.idm.oclc.org/10.1126/science.1176333

Vikramdeo, K.S., Anand, S., Khan, M.A. et al. (2022). Detection of mitochondrial DNA mutations in circulating mitochondria-originated extracellular vesicles for potential diagnostic applications in pancreatic adenocarcinoma. Sci Rep 12, 18455. 10.1038/s41598-022-22006-5

Vogel, R., Pal, A.K., Jambhrunkar, S., Patel, P., Thakur, S.S., Reátegui, E., Parekh, H.S., Saá, P., Stassinopoulos, A., & Broom, M.F. (2017). High-resolution single particle zeta potential characterisation of biological nanoparticles using tunable resistive pulse sensing. Scientific Reports, 7, 17479. 10.1038/s41598-017-14981-x

Wang F, Feng J, Jin A, Shao Y, Shen M, Ma J, Lei L, Liu L. (2025). Extracellular Vesicles for Disease Treatment. Int J Nanomedicine. Mar 17;20:3303–3337. doi: 10.2147/IJN.S506456.vik

Welsh, J.A., Goberdhan, D.C., O’Driscoll, L., Théry, C., Witwer, K.W., et al. (2024). Minimal information for studies of extracellular vesicles (MISEV2023): From basic to advance approaches. Journal of Extracellular Vesicles, 13(2), e12404. 10.1002/jev2.12404

Yanatori, I., Richardson, D.R., Dhekne, H.S., Toyokuni, S., & Kishi, F. (2021). CD63 is regulated by iron via the IRE-IRP system and is important for ferritin secretion by extracellular vesicles. Blood, 138(16), 1490–1503. 10.1182/blood.2021010995

Zhou, Y., Chen, F., Xie, X., Nie, H., Lian, S., Zhong, C., Fu, C., Shen, W., Li, B., Ye, Y., Lu, Y., & Jia, L. (2021). Tumor-derived exosome promotes metastasis via altering its phenotype and inclusions. Journal of Cancer, 12(14), 4240–4246. 10.7150/jca.48043

Xu K, Liu Q, Wu K, Liu L, Zhao M, Yang H, Wang X, Wang W. (2020). Extracellular vesicles as potential biomarkers and therapeutic approaches in autoimmune diseases. J Transl Med. 2020 Nov 12;18(1):432. doi: 10.1186/s12967-020-02609-0.

Xu, S., Jiang, J., Chang, L. et al. Multisystem clinicopathologic and genetic analysis of MELAS. Orphanet J Rare Dis 19, 487 (2024). 10.1186/s13023-024-03511-4

